# Excitatory amino acid transporter 1 supports adult hippocampal neural stem cell self-renewal

**DOI:** 10.1101/2022.10.31.514562

**Authors:** Joshua D. Rieskamp, Ileanexis Rosado-Burgos, Jacob E. Christofi, Eliza Ansar, Dalia Einstein, Ashley E. Walters, Valentina Valentini, John P. Bruno, Elizabeth D. Kirby

## Abstract

Within the adult mammalian dentate gyrus (DG) of the hippocampus, glutamate stimulates neural stem cell (NSC) self-renewing proliferation, providing a link between adult neurogenesis and local circuit activity. Here, we show that glutamate-induced self-renewal of adult DG NSCs requires glutamate transport via excitatory amino acid transporter 1 (EAAT1) to stimulate lipogenesis. Loss of EAAT1 prevented glutamate-induced self-renewing proliferation of NSCs in vitro and in vivo, with little role evident for canonical glutamate receptors. Transcriptomics and further pathway manipulation revealed that glutamate simulation of NSCs relied on EAAT1 transport-stimulated lipogenesis. Our findings demonstrate a critical, direct role for EAAT1 in stimulating NSCs to support neurogenesis in adulthood, thereby providing insights into a non-canonical mechanism by which NSCs sense and respond their niche.

## Introduction

The dentate gyrus of the hippocampus (DG) is one of a few, isolated brain regions that can support resident neural stem cells (NSCs) that generate new neurons in adult mammals. Adult hippocampal NSCs are distinct from developmental counterparts^1–3^, and their persistence through adulthood is conserved across most land-born mammals, likely including primates^4–7^ (but see:^8,9^). The excitatory neurotransmitter glutamate is a potent stimulator of NSC proliferation in the adult DG, a link which suggests a mechanism for tethering adult neurogenesis to circuit activity, as well as a potential mechanism for regeneration in excitotoxic conditions^10,11^.

Recent work shows that glutamate promotes asymmetric, self-renewing NSC divisions^12–14^. However, the molecular mediator by which glutamate stimulates NSCs is unclear. Early studies found that NDMA receptor antagonists actually increase NSC proliferation, rather than decrease it^15–18^. More recently, it was shown that triple transgenic knockout of AMPA, NMDA and metabotropic glutamate receptors in NSCs had no effect on their proliferation^14^. Together, previous work suggests that the NSC response to glutamate is unlikely to rely on simple receptor stimulation.

A major component of glutamate regulation in the adult mammalian brain is the family of excitatory amino acid transporters (EAATs). EAATs are transmembrane proteins that shuttle glutamate from the extracellular to intracellular environment via ion-coupled transport. Robust EAAT expression, in particular EAAT1, is a widely-used marker of adult NSC phenotype^19,20^ and in vivo transport-associated currents suggest these transporters are present and active in the adult DG NSC membrane^21–23^. The functional role of EAATs in NSC regulation has received little attention to date. To better understand how adult NSCs interact with the glutamatergic environment of the adult mammalian DG, we set out to uncover the functional role of EAATs in these cells.

## Results

### High expression of EAAT1, but not other EAATs, is widespread in NSCs and declines rapidly with differentiation

The EAAT family of glutamate transporters has 5 primary members, EAAT1, 2, 3, 4 and 5 (gene names Slc1a3, Slc1a2, Slc1a1, Slc1a6, Slc1a7). Adult DG NSCs have been reported to express both EAAT1 and EAAT2^21^. Using single cell RNAseq (scRNAseq) data from Shin et al.^24^, we confirmed expression of EAAT1 (Slc1a3) and EAAT2 (Slc1a2) in NSCs, with EAAT2 transcript levels being ∼5.35x lower than EAAT1 transcripts (Fig 1A). Both EAAT1 and EAAT2 expression declined significantly with differentiation (pseudotime) (Fig 1B). EAAT3 (Slc1a1), EAAT4 (Slc1a6) and EAAT5 (Slc1a7) were either completely or almost completely undetected.

**Figure 1:**
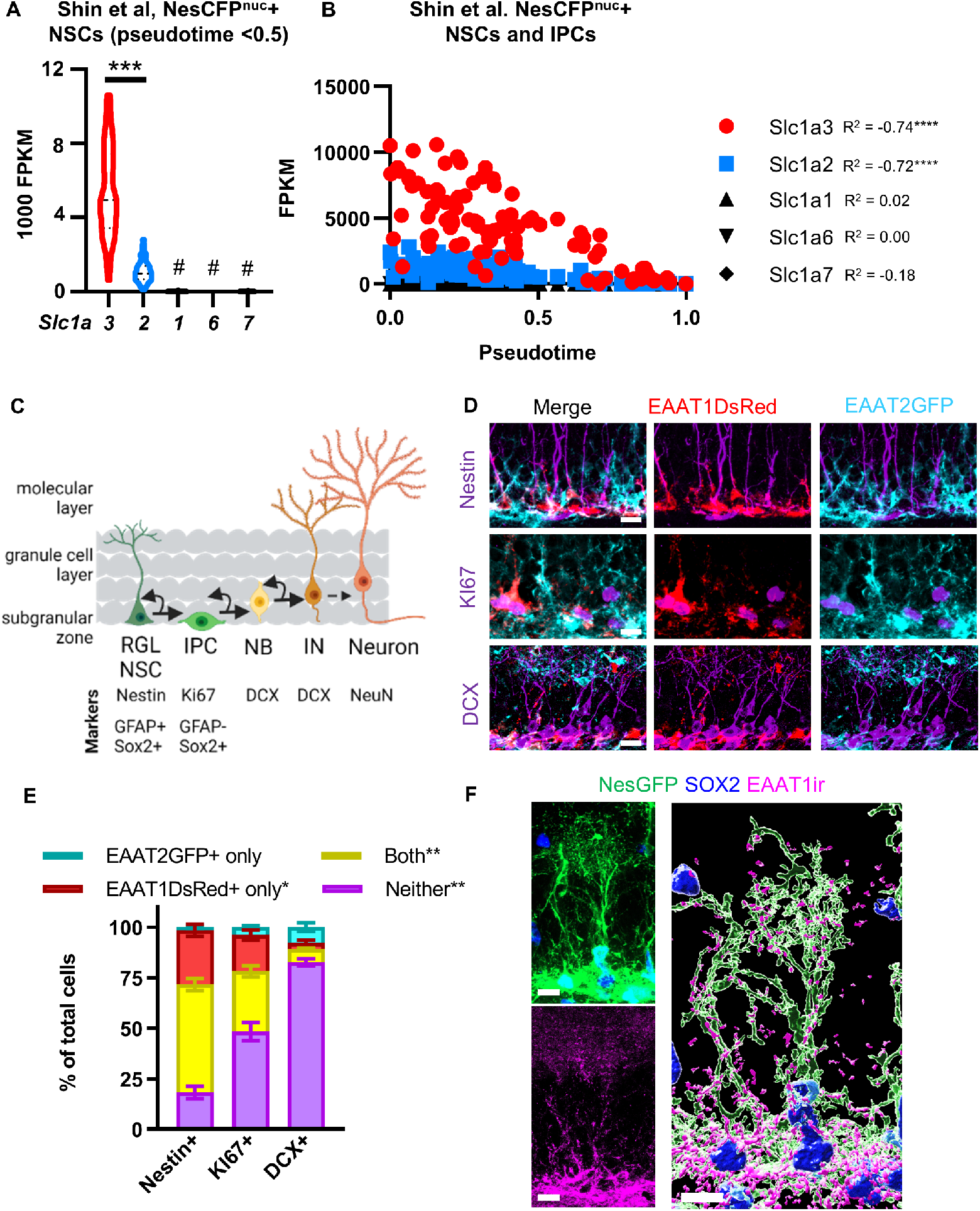
NSC express EAAT1 and to a lesser extent EAAT2. A) scRNAseq FPKM values for EAAT transcripts from data in Shin et al., 2015. Only cells with psuedotime <0.5 included to reflect NSCs (as opposed to early IPCs). ***p<0.001, #p<0.0001 vs EAAT1 and 2, Dunn’s multiple comparisons B) FPKM of EAAT1-5 by pseudotime from Shin et al., 2015. ****p<0.0001 Pearson correlation with pseudotime C) Cells in the neurogenic cascade along with immunophenotypical markers used to identify radial glial like NSCs (RGL NSC), intermediate progenitor cells (IPCs), neuroblasts (NB), immature neurons (IN) and mature neurons. D) Immunolabeling for Nestin, Ki67 and DCX in DG from EAAT1DsRed+EAAT2GFP+ adult mice. E) Proportion of cells expressing EAAT1DsRed+EAAT2GFP+ in adult DG. *p<0.05,**p<0.01 Tukey’s posthoc test. Mean ± SEM of N = 4-5 mice. F) EAAT1 immunolabeling in NestinGFP+ NSC, with Sox2+ nucleus. Merged z-stack on left, 3D reconstruction of z-stack on right. All scales = 10 µm. See also Fig S1.

To verify the extent of EAAT1 and 2 expression throughout the potentially heterogenous population of NSCs in vivo, we used immunolabeling for phenotypic markers in adult EAAT1dsRed and EAAT2GFP transcriptional reporter mice^21^(Fig 1C). 53% of Nestin+ radial glia-like NSCs were positive for both EAAT1dsRed and EAAT2GFP, with another 27% positive for EAAT1dsRed alone (Fig 1D,E, S1A,B). Less than 2% of Nestin+ NSCs were positive for EAAT2GFP alone, leaving 20% of Nestin+ NSCs negative for both transporters. Ki67+ proliferative cells in the subgranular zone, which are primarily composed of IPCs, and doublecortin (DCX+) neuroblasts and immature neurons (NB/IN) each showed progressively smaller proportions of cells expressing EAATs.

To visualize EAAT1 protein distribution throughout the full morphology of NSCs, including their apical process and bushy terminals in the molecular layer, we used EAAT1 immunolabeling in adult NestinGFP mice^25^, as well as with nestin immunolabeling in EAAT1dsRed mice. EAAT1 immunolabeling closely mimicked the distribution of EAAT1dsRed (Fig S1C), with the most intense labeling the subgranular zone. 3D reconstructions also revealed EAAT1 protein putatively co-localizing with the molecular layer bushy terminals of NSCs in both sets of mice (Fig 1F, S1C).

Together, these data suggest that EAAT1 is prominently expressed across the population of adult DG NSCs and EAAT1 protein is distributed throughout the cell, likely including in the apical terminals where glutamatergic synapses are abundant nearby. EAAT2 is also present in NSCs but at lower levels than EAAT1. The data also suggest that expression of both transporters declines rapidly with differentiation into IPC and early neuronal phenotypes.

### Glutamate stimulates adult NSCs via EAAT-mediated transport

To begin investigating the role of EAATs in NSCs, we used cultured NSCs derived from adult male and female DG^26^. These cultures are composed primarily of quiescent and cycling NSCs when maintained in standard conditions^27^. Using our published bulk RNAseq and liquid chromatography tandem mass spectrometry data from these cells^27^, we first confirmed that cultured NSC expression of glutamate transporter and receptor genes was similar to that of in vivo NSCs, with EAAT1 being most abundant, followed by EAAT2, then lower but detectable expression levels of AMPA receptor subunits, NMDA receptor subunits and metabotropic glutamate receptor transcripts (Fig S2A-C). To examine expression of these transporters and receptors throughout the quiescence-activation cycle in cultured NSCs, we used single cell RNAseq transcriptome data from^27^. EAAT1 (Slc1a3) and AMPA/mGluR receptor genes were expressed most abundantly in quiescent and cycling NSCs, with downregulation generally evident in those exiting the cell cycle (presumably to differentiate) (Fig S2D). These findings generally agree with in vivo NSC scRNAseq (Fig 1B), showing high Slc1a3 in quiescent/active NSCs (pseudotime < ∼ 0.5) and downregulation with differentiation. Together, these data show that cultured NSCs have similar expression patterns of glutamate transporters and receptors as in vivo NSCs.

We next confirmed that glutamate stimulates proliferation of isolated NSCs using incorporation of the thymidine analog ethynyl deoxyurdine (EdU) after glutamate exposure. Glutamate increased NSC proliferation ∼2-fold after 48h of treatment, with the effect being most notable with 100 µM glutamate (Fig 2A,B). To determine the role of glutamate receptors versus transporters in this glutamate-induced proliferation, we treated NSCs with several pharmacologic inhibitors. Treatment with a cocktail of inhibitors to block AMPA, NMDA and mGluR5 (NBQX, D-AP5, and ACDPP, referred to together as αGlut R) had no effect on basal NSC proliferation nor their response to glutamate (Fig 2C,D). Treatment with the panEAAT inhibitor TFB-TBOA (αpanEAAT), in contrast, blocked glutamate-induced proliferation. Transport inhibition had this effect whether glutamate receptors were blocked or not. EAAT1 and 2 both transport aspartate, as well as glutamate. We repeated the experiment with aspartate and found that, similar to glutamate, aspartate stimulated NSC proliferation and that effect was blocked by αpanEAAT inhibition (Fig 2E,F). Aspartate can activate NDMA receptors at higher doses. Including the NMDA receptor inhibitor (D-AP5) had no effect on NSC proliferation in any condition. These findings suggested, unexpectedly, that EAAT transporter activity alone mediates glutamate-induced NSC proliferation.

**Figure 2:**
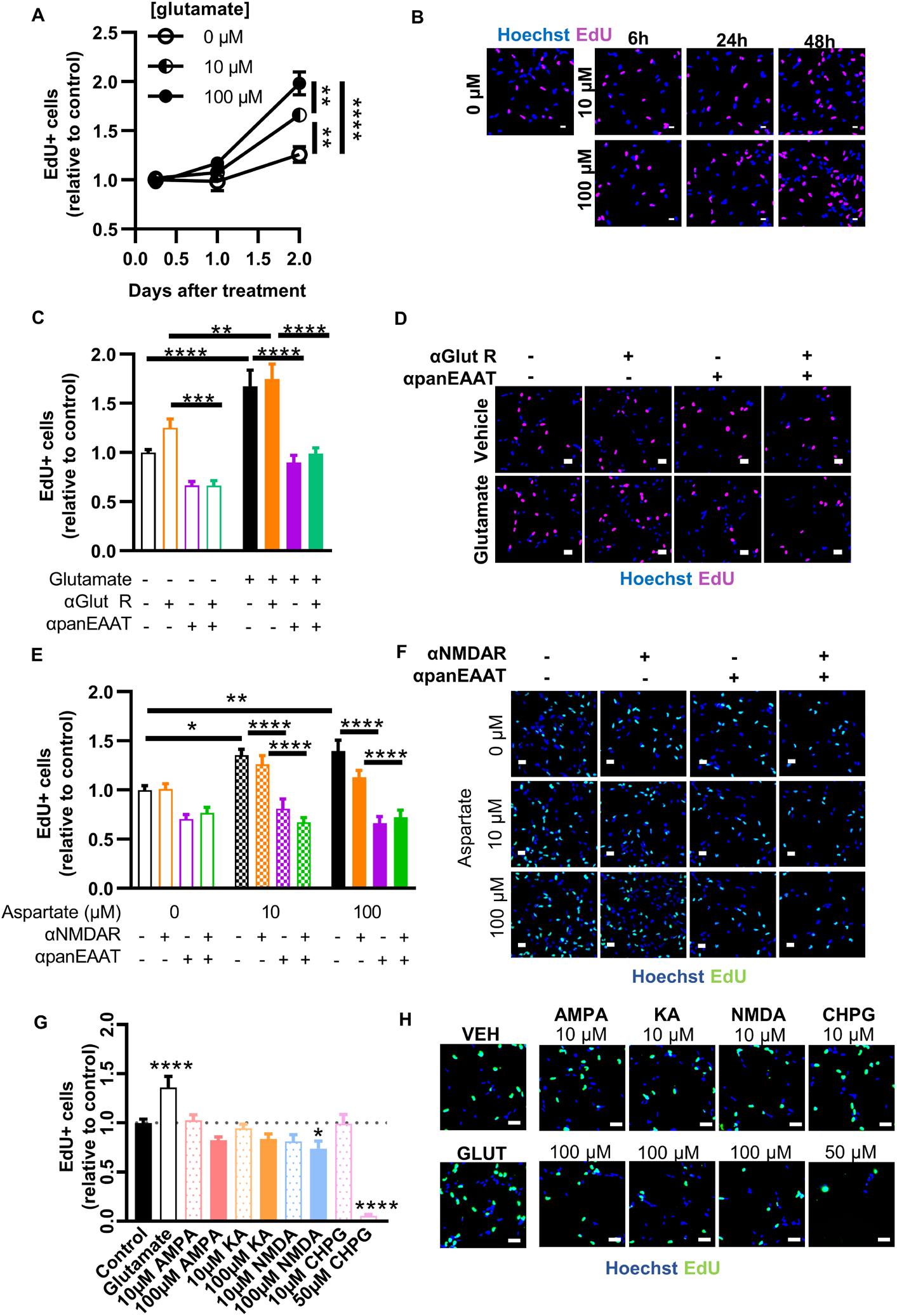
Glutamate stimulates NSC proliferation via EAATs. A) EdU+ cell counts after glutamate treatment over time. **p<0.01, ****p<0.001 Tukey’s comparisons within timepoint. Mean ± SEM of N = 2-5 replicates/experiment, 3 independent experiments. B) Representative image of EdU+ labeling in cultured NSCs after glutamate treatment C) EdU+ cell counts after treatment with glutamate +/- inhibitors of glutamate receptors (αGlutR, NBQX, D-AP5, and ACDPP) or EAATs (αpanEAAT, TFB-TBOA). **,***,****p<0.01, 0.001, 0.0001 Sidak’s multiple comparisons. Mean ± SEM of N = 5 replicates/experiment, 4 independent experiments. D) Representative image of EdU+ labeling in cultured NSCs after glutamate +/− receptor and transporter inhibitors E) EdU+ cell counts after treatment with aspartate +/− inhibitor of NMDA receptor (αNMDAR) or EAATs (αpanEAAT). *,**,***,****p<0.05, 0.001, 0.0001 Tukey’s multiple comparisons. Mean ± SEM of N = 5 replicates/experiment, 3 independent experiments. F) Representative image of EdU+ labeling in cultured NSCs after aspartate treatment. G) EdU+ cell counts after treatment with glutamatergic agonists. *,****p<0.01, 0.0001 Dunnett’s multiple comparisons to control. Mean ± SEM of N = 6-12 replicates/experiment, 4 independent experiments. H) Representative image of EdU+ labeling in cultured NSCs after glutamate receptor agonist treatment. Hoechst to label nuclei and scale = 20 µm throughout. See also Fig S2

To further explore the role of glutamate receptors in NSC proliferation, we treated NSCs with glutamate receptor agonists: AMPA, KA, NMDA and CHPG (mGluR5 agonist). None of the agonists increased proliferation (Fig G,H). NMDA slightly decreased proliferation. CHPG (mGluR5) agonist caused massive cell death at its higher dose and had no effect at its lower dose. These findings suggest, unexpectedly, that EAAT transporter activity alone mediates glutamate-induced NSC proliferation with no, or possibly a countering, contribution of glutamate receptors.

### Glutamate stimulation of adult NSCs relies on EAAT1 not EAAT2

To determine which EAATs might be active in NSCs, we measured glutamate clearance from NSC cultures using ultra-high performance liquid chromatography with electrochemical detection (uHPLC-ECD) of conditioned media. Within 20 minutes, NSCs cleared 82% of a 5 µM glutamate pulse from the media (Fig 3A). Treatment with the selective EAAT1 inhibitor (UCPH101, αEAAT1) or a combination of αEAAT1 and the selective EAAT2 inhibitor (WAY213613, αEAAT2) completely prevented glutamate clearance compared to vehicle treated controls (Fig 3B). αEAAT2 alone had no effect on clearance compared to vehicle treated controls. To confirm activity of the αEAAT2 drug, we replicated the experiment with a higher dose that also inhibits EAAT1. Higher dose αEAAT2 partially prevented glutamate clearance, suggesting that the compound is active but only shows effects at doses that inhibit EAAT1 (Fig S3A). Similar results were found with an alternative EAAT2 inhibitor (DHK) (Fig S3B). Taken together, these results demonstrate functional glutamate clearance by NSCs and implicate EAAT1 as the dominant transporter.

**Figure 3:**
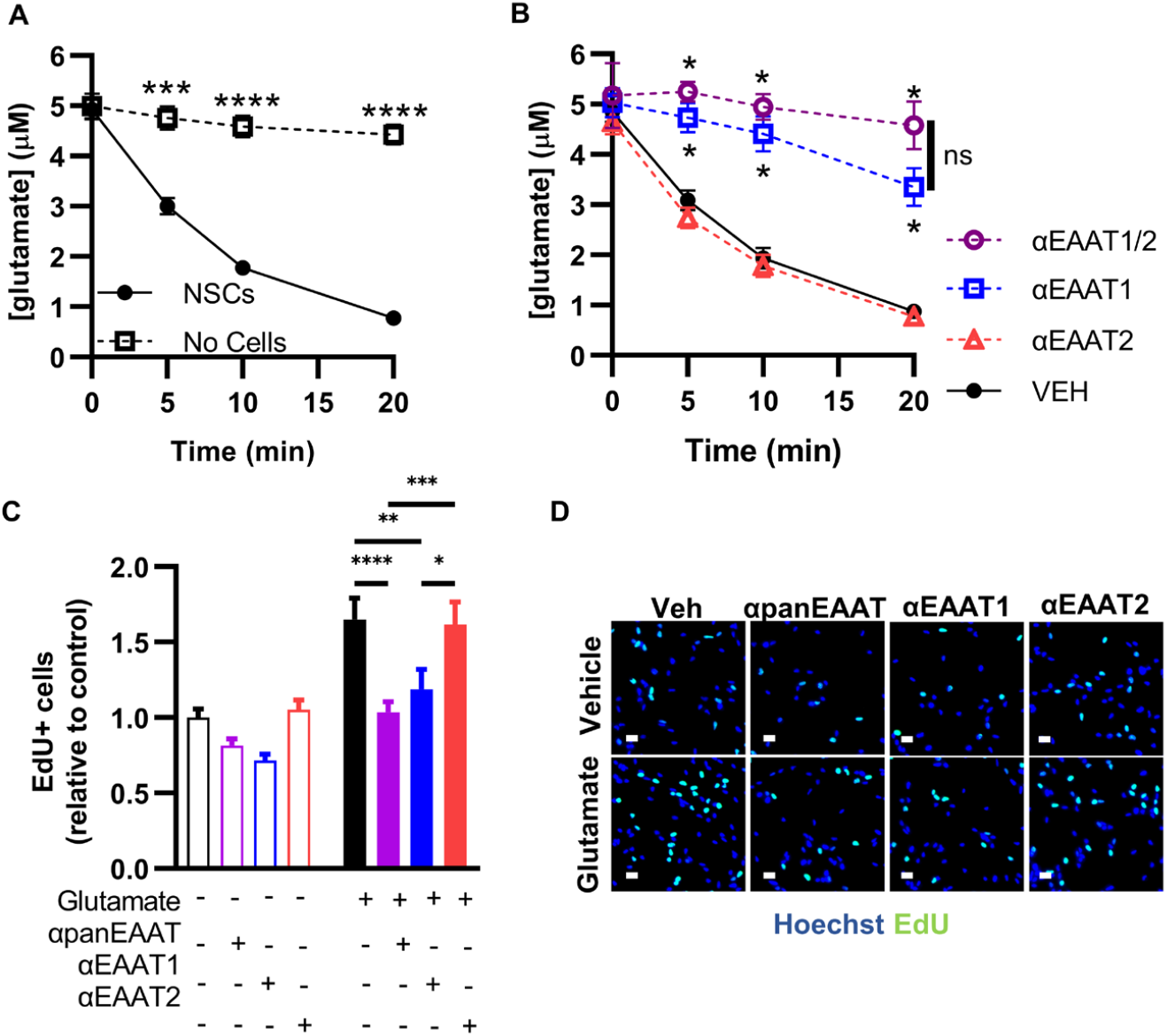
Glutamate stimulates NSC proliferation via EAAT1. A) Glutamate in conditioned media as measured by uHPLC after a 5 µM pulse in wells with NSCs or empty wells. ***,****p<0.001, 0.0001 vs “No Cells” Sidak’s multiple comparisons. Mean ± SEM of N = 6 independent experiments. B) Glutamate in NSC conditioned media as measured by uHPLC after a 5 µM pulse with pretreatment with inhibitors of EAAT1 (UCPH102), EAAT2 (WAY213613) or their combination (EAAT1/2). *p<0.05 Tukey’s comparisons within timepoint. Mean ± SEM of N = 3 independent experiments. C) EdU+ cell counts after treatment with glutamate +/− inhibitors of EAAT1 (UCPH102), EAAT2 (WAY213613), or panEAAT (TFB-TBOA). *,**p<0.05, 0.01 Tukey’s multiple comparisons. Mean ± SEM of N = 5 replicates/experiment, 3 independent experiments. EdU+ counts expressed relative to 0 µM glutamate/vehicle within experiment. D) Representative image of EdU+ labeling in cultured NSCs after glutamate +/− transporter inhibitors. Hoechst to label nuclei and scale = 20 µm. See also Fig S3.

To test the functional role of EAAT1 vs EAAT2 in NSC proliferation, we treated cultured NSCs with αEAAT1, αEAAT2 or αpanEAAT then quantified cell proliferation using EdU labeling. Inhibiting EAAT1 alone or panEAAT inhibition prevented glutamate-induced NSC proliferation while EAAT2 inhibition alone had no effect (Fig 3C,D). Throughout these experiments, cells in all groups appeared healthy and showed no observable signs of apoptosis (fragmented nuclei, loss of plate adhesion). We also confirmed that neither αpanEAAT nor αEAAT1 treatment led to signs of cell death using a Sapphire700 IR dye assay (Fig S3C). Together, these data suggest that glutamate stimulates NSC proliferation directly via EAAT1 activity.

### EAAT1 knockdown causes cell-autonomous loss of NSCs in vivo

To examine the role of EAAT1 in DG NSCs in vivo, we used CRISPR interference (CRISPRi), a transcriptional interference strategy based on the CRISPR/cas9 system. In CRISPRi, a catalytically dead Cas9 protein (dCas9) fused to a Kruppel associated box (KRAB) domain represses transcription of a gene targeted by a highly specific single guide (sg)RNA. We designed a U6àsgRNA against EAAT1 (EAAT1 KD) (and a non-target sgRNA (NT)) then inserted it into a lentiviral CRISPRi backbone^28^ with dCas9-KRAB-T2A-GFP (Fig 4A). Cultured NSCs infected with the EAAT1 KD CRISPRi vector showed a 53.1% loss of EAAT1 immunoreactivity and significant suppression of proliferation compared to NT infected cells (Fig S4A-D). We stereotaxically infused the EAAT1 KD and NT CRISPRi lentiviral vectors in the DG of adult mice (Fig 4A,B). 1 week later, no loss of EAAT1 protein was detected (Fig S3E,F). 3 weeks later, a 40.6% knockdown of EAAT1 immunoreactivity was detected in EAAT1 KD compared to NT treated controls (Fig 4C,D). We therefore next assessed the effect of EAAT1 loss on the neurogenic cascade at this 3 week timepoint.

**Figure 4:**
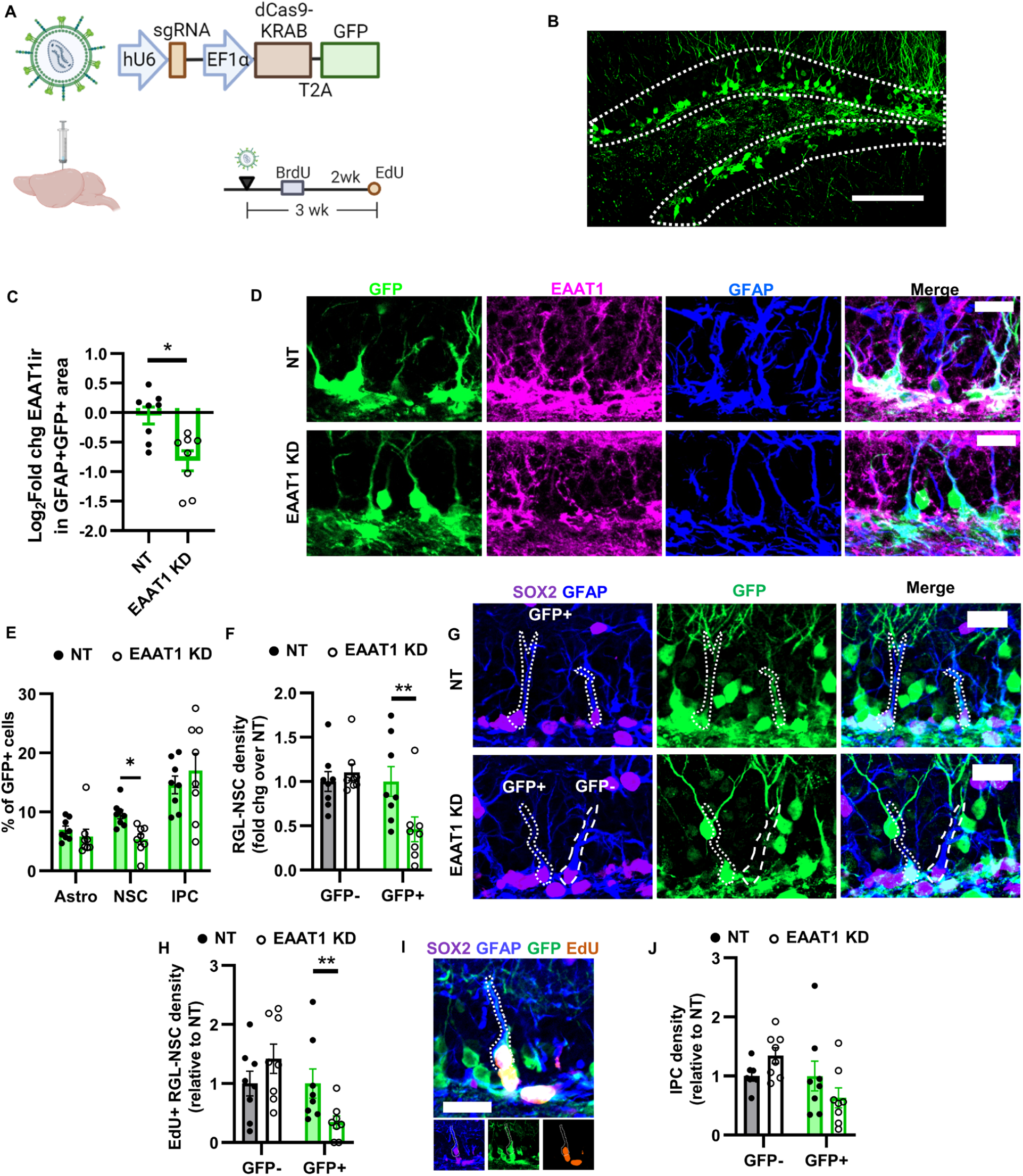
EAAT1 knockdown causes cell-autonomous loss of the NSC pool. A) CRISPRI lentiviral vector design and timeline. B) Representative GFP+ labeling in DG at 3 weeks after infusion. Scale = 100 µm. C) EAAT1 immunolabeling intensity within GFP+GFAP+ area of mice infused with NT or EAAT1 KD CRISPRi lentiviral vectors in the DG. D) Representative image of EAAT1 labeling coupled with GFP and GFAP at 3 weeks after CRISPRi infusion. Co-labeling becomes white in merged image. Scale = 20 µm. E) Percent of GFP+ cells expressing phenotypic markers of astrocytes (GFAP+SOX2+ stellate), NSCs (GFAP+SOX2+ radial) or IPCs (GFAP-SOX2+) 3 weeks after CRISPRi infusion. F) Density of GFP+ and GFP-GFAP+SOX2+ radial glia-like NSCs in NT and EAAT1 KD mice 3 weeks after infusion. G) Representative images of GFAP, SOX2, GFP immunolabeling 3 weeks after infusion. GFP+ and GFP-NSCs are outlined. Scale = 20 µm. H) Density of GFP+ and GFP-Edll+GFAP+SOX2+ proliferating radial glia-like NSCs in NT and EAAT1 KD mice 3 weeks after infusion I) Representative image of an EdU+GFP+GFAP+SOX2+ radial glia-like NSC 3 weeks after infusion. Scale = 10 µm. J) Density of GFP+ and GFP-GFAP-SOX2+ IPCs in NT and EAAT1 KD mice 3 weeks after infusion. All densities relative to NT control within GFP type. Mean ± SEM of N = 8 individual mice (points shown). *,**p<0.05, 0.01 Sidak’s multiple comparisons, except panel C in which *p<0.05 Mann Whitney test. See also Fig S4

At 3 weeks after CRISPRi infusion, ∼30% of dCas9-KRAB-GFP+ NSCs (hereafter GFP+ NSCs) were SOX2+ astrocytes, NSCs or IPCs, the cell types which express EAAT1 (Fig S4G). Among those, we found a selective reduction in the percent of GFP+ NSCs, but not in the percent of GFP+ astrocytes or IPCs (Fig 4E). In this paradigm, EAAT1 knockdown could alter NSCs via receptor-independent, cell-autonomous loss of glutamate transport, as our in vitro data suggests, or through increased glutamate stimulation of receptors due to extracellular glutamate buildup, or through a combination of the two. While changes in ambient glutamate would affect both GFP+ and GFP-NSCs, only the GFP+ NSCs would be lacking cell autonomous glutamate transport. To investigate the cell autonomous effect, we therefore compared GFP+ NSCs to neighboring GFP-NSCs. At 3 weeks after viral infusion, EAAT1 KD mice had 53.9% fewer GFP+GFAP+SOX2+ radial glia-like NSCs compared to NT treated controls (Fig 4F,G). In contrast, the neighboring GFP-NSCs in these mice showed no effect of EAAT1 KD. The number of proliferating (EdU+) GFP+GFAP+ radial glia-like NSCs showed the same pattern, with a 63.5% decrease in density of proliferating GFP+ NSCs with EAAT1 KD but no effect on neighboring GFP-proliferating NSCs compared to NT control mice (Fig 4H,I). No differences were found between NT and EAAT1 KD mice in GFP+ or GFP-GFAP-SOX2+ IPC density nor in density of DCX+ neuroblasts/immature neurons that were BrdU-labeled 2 weeks before perfusion (Fig 4J, S4H,I). These findings suggest that loss of EAAT1 causes a cell-autonomous loss of the radial glia-like NSC pool.

### Long-term EAAT1 knockdown causes cell-autonomous loss of adult neurogenesis

To determine whether the loss of NSCs observed at 3 weeks after knockdown would result in loss of adult neurogenesis at longer timepoints, we perfused mice 2 months after NT or EAAT1 KD CRISPRi lentiviral infusion (Fig 5A). The cell autonomous loss of GFP+ NSCs persisted at this later timepoint. EAAT1 KD mice showed a 63.0% loss of GFP+ radial glia-like NSCs with no effect on GFP-NSCs compared to NT control mice (Fig 5B). GFP+ IPC density was also 54.3% lower in EAAT1 KD mice compared to NT control mice, while density of GFP-IPCs did not differ between EAAT1 KD and NT mice (Fig 5C). EAAT1 KD mice also showed a 73.6% reduction in number of GFP+NeuN+ newborn neurons labeled with BrdU 1 month before perfusion (Fig 5E,F). These data suggest that loss of EAAT1 results in loss of adult neurogenesis that is restricted to the progeny of EAAT1 KD NSCs and does not extend to neighboring NSCs and their progeny.

**Figure 5:**
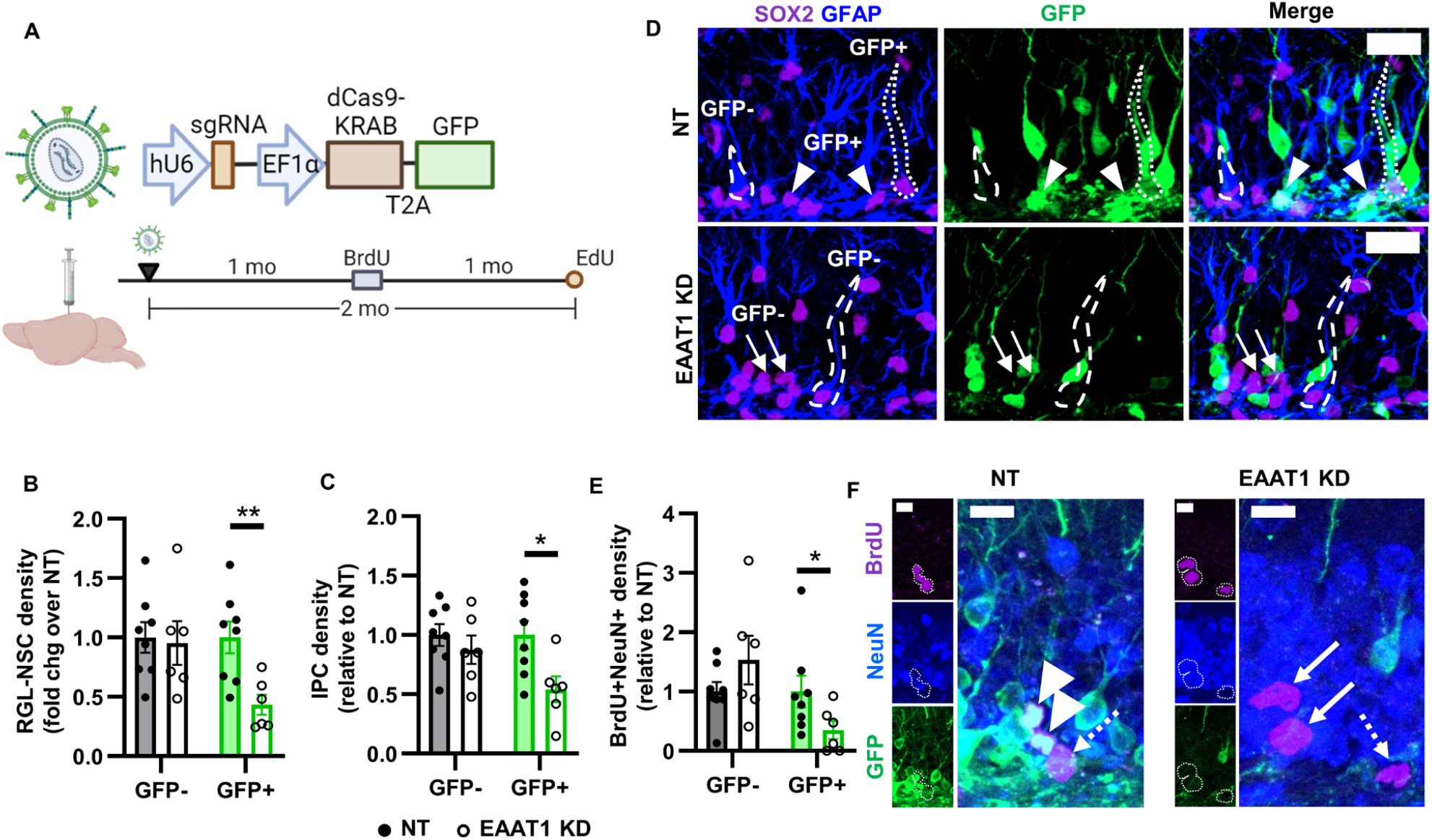
EAAT1 knockdown causes cell-autonomous loss of neuronal NSC progeny. A) Timeline for CRISPRi infusion and thymidine analog administration. B) Density of GFP+ and GFP-GFAP+SOX2+ radial glia-like NSCs in NT and EAAT1 KD mice 2 months after infusion. C) Density of GFP+ and GFP-GFAP-SOX2+ IPCs in NT and EAAT1 KD mice 2 months after infusion. D) Representative images of GFAP, SOX2, GFP immunolabeling 2 month after infusion. GFP+ and GFP-NSCs are outlined. IPCs shown by arrows (GFP-) or arrowheads (GFP+). Scale = 20 µm. E) Density of GFP+ and GFP-NeuN+BrdU+ cells in NT and EAAT1 KD mice 2 months after infusion. F) Representative images of BrdU+ cells. Arrowhead = BrdU+NeuN+GFP+, Arrow = NeuN+Brdll+, Dashed arrow = BrdU+. Scale = 10 µm All densities relative to NT control within GFP type. Mean ± SEM of N = 6-8 individual mice (points shown). *,**p<0.05, 0.01 Sidak’s multiple comparisons within GFP label type

### EAAT-dependent stimulation of fatty acid synthetase is essential for glutamate-induced NSC proliferation

To gain insight into the molecular mediators of EAAT-dependent stimulation of NSC self-renewal, we performed bulk RNAseq on RNA from cultured NSCs treated with glutamate and/or panEAAT inhibitor. Principle component analysis revealed a strong separation of groups by EAAT inhibition status (Fig 6A). In the absence of αpanEAAT, glutamate treatment caused significant upregulation of 106 transcripts and significant downregulation of 49 transcripts (Table S1). In contrast, in the presence of αpanEAAT, glutamate treatment yielded no upregulated transcripts and only 1 downregulated transcript. These data suggest robust NSC transcriptional response to glutamate that is prevented by EAAT inhibition.

**Figure 6:**
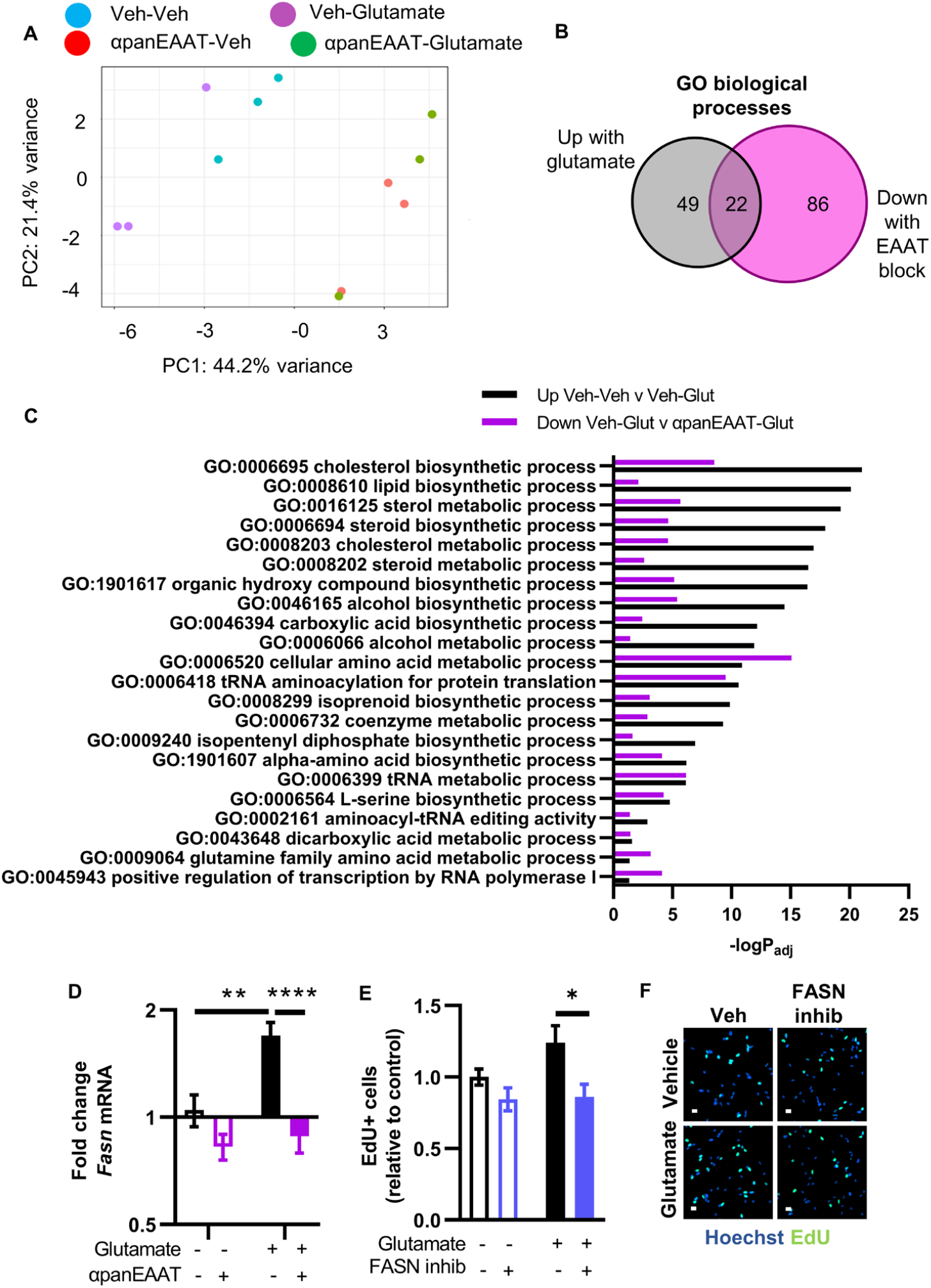
Glutamate stimulates NSC self-renewal via transport-stimulated lipogenesis. A) Principle component analysis of RNAseq samples from treated in vitro NSCs. B,C) 22 GO biological processes that were upregulated by glutamate and downregulated panEAAT inhibition (TFBTBOA) in RNAseq. -logPadj shown. N = 3 replicates/group. D) Fasn mRNA expression as measured by real time qPCR after treatment with glutamate +/− panEAAT inhibition (TFBTBOA). **,****p<0.01, 0.0001 Sidak’s multiple comparisons. Mean ± SEM of N = 3 replicates/experiment, 3 independent experiments. E) EdU+ cell counts after treatment with glutamate +/− inhibitor of FASN (orlistat). Mean ± SEM of N = 4 replicates/experiment, 4 independent experiments. *p<0.05 Sidak’s multiple comparisons. F) Representative images of EdU labeling in NSCs treated with glutamate +/- FASN inhibitor orlistat. Hoechst as nuclear marker. Scale = 10 µm.

To understand the functional significance of genes regulated by glutamate transport, we focused on GO biological process analysis of genes upregulated by glutamate that were also downregulated by αpanEAAT (Table S2). We found 22 overlapping GO terms which centered mostly on metabolic processes related to lipid, amino acid, and protein synthesis (Fig 6B,C).

Particularly notable among the cellular processes implicated in EAAT-dependent regulation of NSC gene expression was lipid biosynthetic process. Lipogenesis is essential for NSC self-renewal, with a particularly critical role for fatty acid synthetase (FASN)^29–31^. In our RNAseq data, glutamate induced upregulation of Fasn, an effect that EAAT inhibition blocked (Table S1,S3). Independent replication using qPCR confirmed that glutamate stimulated Fasn expression in an EAAT-dependent manner (Fig 6D). To test the functional role of Fasn in glutamate-stimulated NSC proliferation, we treated cultured NSCs with the FASN inhibitor, orlistat. FASN inhibition prevented glutamate-induced NSC proliferation (Fig 6E,F). These findings suggest that glutamate transport by EAAT promotes NSC proliferation via stimulation of FASN.

## Discussion

Our findings suggest a functional role for EAAT1 in supporting adult hippocampal neurogenesis via cell autonomous NSC pool maintenance. Our findings stand in contrast to a canonical glutamate receptor-dependent mechanism reported in most other cell types and suggest that EAAT1 expressed by NSCs directly supports self-renewing proliferation via transport-dependent stimulation of lipogenesis.

Our findings showed that loss of EAAT1 led to cell-autonomous loss of radial glial-like NSCs and their progeny in a pattern that suggests impaired asymmetric, self-renewing division. The majority of adult NSCs are quiescent and the maintenance of that quiescence is essential for sustaining adult neurogenesis throughout the lifespan^32,33^. Ectopic activation from quiescence can result in exhaustion of the NSC pool but it is typically preceded by a surge in NSC proliferation^34–36^. We did not observe any increase in NSC proliferation after EAAT1 knockdown, suggesting that quiescence itself is not regulated by EAAT1. Rather, our data suggest a shift in cell division mode that impairs self-renewal. When NSCs activate, they can either undergo asymmetric or symmetric divisions. Asymmetric (self-renewing) division results primarily in production of an NSC and IPC. Symmetric division can take either expansive (two NSCs) or exhaustive (two IPCs) form. Exhaustive, symmetric division depletes the NSC pool and reduces neurogenesis. Because we observed a progressive loss of total and proliferating NSCs that over longer time periods expanded to the IPC and new neuron populations, our data suggest that EAAT1 cell-autonomously supports self-renewing cell division. EAAT1 loss therefore putatively results in a shift to exhaustive divisions that deplete the neurogenic pool.

Expression of glutamate transporter proteins, particularly EAAT1, is widespread in the radial glia NSCs of the embryonic forebrain^37^. One study in early postnatal DG NSCs concluded that EAATs indirectly suppress proliferation by limiting the activation of mGluRs^38^. This apparent reversal of the roles of EAAT1 and mGluR5 in early postnatal hippocampus versus what we observed in adult DG NSCs is paralleled by downregulated expression of mGluR5 and upregulated expression of EAAT1 in DG NSCs during the transition to adulthood^39^. This transition could contribute to the shift in cellular behavior of DG NSCs from rapidly proliferating and depleting early in life to more maintenance-focused later in life^33^. Comparative studies of EAAT function in NSCs across different stages of development could shed further light on this hypothesis.

Cell metabolic states are becoming increasingly recognized as active players in NSC self-renewal and fate decisions, both as generators of key cellular substrates for growth and as independent initiators of gene expression^40–42^. Several previous studies demonstrate a particularly critical role for fatty acid synthesis in DG NSC self-renewal^29–31^. In agreement with this previous data, we found that FASN activity was required for glutamate-induced NSC proliferation. Cell metabolism, and lipogenesis in particular, therefore represent a potential convergence point for the glutamate-stimulated, EAAT-dependent processes that we observed in adult DG NSCs. A remaining open question, however, is how transport of glutamate stimulates lipogenesis. Glutamate transport both provides glutamate as a substrate to the intracellular environment and a net inward positive current due to ionic co-transport^43^. Either or both of these stimuli could underpin the connection between glutamate transport and lipogenesis. Future research will be needed to address these questions.

NSCs may derive glutamatergic input from multiple sources in vivo. NSC apical processes wrap around putatively glutamatergic synaptic terminals in the molecular layer of the DG^44^. Inputs here from hilar mossy cells^14^ and hypothalamus^13^ both stimulate NSC self-renewal, as does release from local astrocytes^12^. With multiple sources of glutamate, it is tempting to speculate that changes in hippocampal glutamate signaling that occur during normal development, neurodegenerative pathology, and plasticity-inducing stimuli^45–48^ could be transduced by EAAT1 into metabolic states that regulate the population maintenance of NSCs. Future studies could examine whether changes in EAAT expression or function modulate the response of NSCs to these physiological states.

In summary, we show that glutamate transport via EAAT1 cell-autonomously supports DG NSC self-renewal in adult mice. These findings help resolve some discrepancy in the field about the role of glutamate receptors versus glutamate itself in NSC maintenance. They also raise additional questions about the interplay between glutamate signaling and NSC behavior in contexts such as biological aging, neurological disorders, and plasticity-inducing stimuli like exercise and enriched environment^10,49,50^.

## Limitations of the study

The presented studies only examine NSCs in the adult DG of mice. The NSCs in the adult subventricular zone neurogenic niche also express EAAT1^51^, and the role of EAAT1 in this niche remains an open question, to the best of our knowledge. Whether these results extend to other species, including primates, is also not yet clear either. However, a recent scRNAseq profiling of adult macaque hippocampal NSCs show high levels of EAAT1 and EAAT2 expression^52^, suggesting the possibility that NSCs in adult primate hippocampus could rely on EAAT-mediated glutamate transport. We also do not resolve which glutamatergic inputs impact NSCs directly via EAAT1, nor how EAATs come in to play in pathological conditions characterized by excess glutamate transmission. Our work also only begins the uncovering of the intracellular signaling between glutamate transport and stimulation NSC self-renewal. These steps will require more detailed investigation in the future.

## Methods

### Animals

All experimental protocols were performed in accordance with institutional guidelines approved by the Ohio State University Institutional Animal Care and Use Committee (and with recommendations of the National Institutes of Health Guide for the Care and Use of Laboratory Animals. For lentiviral infusion, seven week old C57BL/6J mice (JAX # 000664) were obtained from The Jackson Laboratory were allowed to acclimate for 1 week prior to surgery. GLAST/EAAT1-DsRed and GLT-1/EAAT2-GFP mice (Regan et al., 2007) were obtained as a kind gift from Dr. Jeffrey Rothstein and crossed to obtain offspring used in experiments. From the time of weaning, mice were group housed (5 per cage) in ventilated cages with ad libitum access to food and water and maintained on a 12 hour light cycle. Reporter mice aged 8 – 12 weeks were used for tissue staining. Both male and female mice were used in all experiments.

### Perfusion and tissue harvest

Mice were injected with a ketamine/xylazine mixture (87.5 mg/kg, 12.5 mg/kg, i.p.) and transcardially perfused with ice-cold PBS. Brains were collected and fixed for 24h in 4% paraformaldehyde in 0.1 M phosphate buffer prior to equilibration for at least 2 days in 30% sucrose in PBS, both at 4°C. They were then sliced on a freezing microtome (Leica) in a 1 in 12 series of 40 µm slices and stored in cryoprotectant at −20°C.

### Immunofluorescence and thymidine analog detection in brain tissue

Sections were rinsed 3x with PBS and then incubated in 1% normal donkey serum (Jackson ImmunoResearch), 0.3% Triton X-100 (blocking buffer) for 1 hour before incubating in primary antibodies (Table 9) diluted in blocking buffer overnight at 4°C with rocking. On the second day, sections were rinsed 3x with PBS and incubated in fluorophore-conjugated secondary antibodies diluted 1:500 in blocking buffer for 2 hours before incubation for 10 min in Hoechst 33342 (Fisher, 1:2000 in PBS), except for sections that were instead incubated with AlexaFluor 350 (Invitrogen) secondary antibodies, and a final 3x PBS rinse. They were then mounted on charged glass slides and coverslipped with ProLong Gold fluorescence mounting media.

For EAAT1 immunolabeling, the above procedure was followed except that tissue was pretreated with ice cold methanol for 10 min for antigen retrieval and incubated in primary antibody at 37°C overnight with shaking to increase antibody penetration.

For nestin immunolabeling, antigen retrieval was performed using citrate buffer, pH = 6.0, 10 min at 95 °C prior to blocking and primary antibody incubation.

For detection of thymidine analogs, the staining process above was used with the following modifications. Tissue sections were incubated for 30 minutes in 2 N HCl at 37°C prior to incubation in anti-BrdU primary antibodies. For detection of EdU, adherent cells or tissue sections were permeabilized for 20 minutes in 0.5% Triton X-100 and incubated in the dark for 30 minutes with copper-dependent click reaction mixture (Click Chemistry Tools #1350) to label EdU with Azide-biotin (Click Chemistry Tools #1265) for subsequent Streptavidin-555 (Invitrogen) conjugation.

Antibody dilutions are listed below:

| Host | Target | Vendor | Cat # | Dilution |
| --- | --- | --- | --- | --- |
| **Rabbit** | mCherry/DsRed | Abcam | ab167453 | 1:500 |
| **Mouse** | mCherry/DsRed | Novus Biologicals | NBP1-96752 | 1:500 |
| **Goat** | GFP | Abcam | ab6673 | 1:1000 |
| **Mouse** | Nestin | Millipore-Sigma | MAB353 | 1:100 |
| **Rat** | KI67 | Invitrogen | 14-5698-82 | 1:500 |
| **Rabbit** | DCX | Cell Signaling | 4604 | 1:500 |
| **Rabbit** | EAAT1 | Abcam | ab416 | 1:250 |
| **Rabbit** | GFAP | Dako | ZO334 | 1:1000 |
| **Rat** | SOX2 | Affymetrix | 14-9811 | 1:1000 |
| **Rat** | BrdU | BioRad | OT0030 | 1:500 |
| **Mouse** | NeuN | Millipore | MAB377 | 1:500 |
| **Rabbit** | EAAT1 | Abcam | Ab416 | 1:500 |

### Image acquisition and analysis

All imaging was performed on a Zeiss Axio Observer Z.1 with apotome digital imaging system and Axiocam 506 monochrome camera (Zeiss). Images of cultured NSCs captured using Zen Blue software (Zeiss) were converted to Tag Image File Format (Tiff) for automated cell counting using ImageJ. Tissue sections were imaged using a 20x objective as z-stacks with 20 × 1 µm steps. Images were analyzed using Zen Blue software (Zeiss). RGLs and IPCs, EdU+ cells, DCX+BrdU+ cells and NeuN+BrdU+ cells were counted and/or phenotyped by colocalization with cell type identity markers within the SGZ (or SGZ plus GCL for BrdU+ cells). Imaging and cell quantification were performed while blind to animal identity. 3D reconstructions were performed using 0.5µm z-stacks taken with a 63x oil objective and then rendered with Imaris software (Oxford Instruments). EAAT1 immunolabeling for quantification of knockdown in brain tissue was done using ImageJ to select GFAP+GFP+ area and measure EAAT1 intensity within that area.

### Cell culture

NSCs were derived from the DG of 6 week old C57BL/6J mice (JAX # 000664) mice based on a published protocol^26^. Briefly, mice were euthanized by CO_2_ inhalation and brains were rapidly dissected. Bilateral DG were isolated from each brain in ice-cold Neurobasal A media (Gibco) and minced with a scalpel blade prior to 20 min incubation in papain (Roche)/dispase (StemCell Technologies)/DNase at 37 °C. The mixture was then triturated with a P1000 pipette and centrifuged for 2 min at 800 x g. The pellet was resuspended in warm Neurobasal A and mixed with a 22% Percoll suspension prior to centrifugation for 15 min at 450 x g. The resulting pellet was washed several times in warm Neurobasal A prior to resuspension in complete growth media containing 1% B27 supplement (Gibco), 0.5% GlutaMax (Gibco), and 20 ng/mL each of epidermal growth factor (EGF) and basic fibroblast growth factor (FGF2) (Peprotech). NSCs were maintained in adherent monolayer culture on poly-D-lysine (Sigma) and laminin (Invitrogen) coated plates. All experiments used cells between passages 4 and 20 and were replicated in two separate cell lines, one derived from 4 pooled male DG and the other from 4 pooled female DG. Both lines were mycoplasma tested and confirmed to produce neurons and glia upon growth factor withdrawal as previously described^27^.

### Pharmacological reagents

Reagents and working dilutions are in Table 1 below:

| Compound | Vendor | Cat # | Working [ ]<br>(μM) | Diluent |
| --- | --- | --- | --- | --- |
| **L-glutamic acid (glutamate)** | Millipore-Sigma | G8415 | 5, 10, 100 | equimolar NaOH |
| **L-aspartic acid (aspartate)** | Tocris | 214 | 10, 100 | equimolar NaOH |
| **TFB-TBOA** | Tocris | 2532 | 1 | DMSO |
| **UCPH-102** | Abcam | ab146404 | 10 | DMSO |
| **Dihydrokainic acid (DHK)** | Tocris | 0111 | 100 | PBS |
| **WAY 213613** | Tocris | 2652 | 1, 10 | DMSO |
| **NBQX disodium salt** | Tocris | 1044 | 10 | PBS |
| **D-AP5** | Tocris | 0106 | 100 | PBS |
| **ACDPP hydrochloride** | Tocris | 2254 | 1 | DMSO |
| **(S)-AMPA** | Tocris | 0254 | 10, 100 | PBS |
| **Kainic acid** | Millipore-Sigma | 420318 | 10, 100 | PBS |
| **NMDA** | Tocris | 0114 | 10, 100 | PBS |
| **CHPG sodium salt** | Tocris | 3695 | 10, 50 | PBS |
| **SEA 0400** | Tocris | 6164 | 1 | DMSO |
| **YM 244769** | Tocris | 4544 | 2 | DMSO |
| **Dantrolene, sodium salt** | Tocris | 0507 | 10 | DMSO |
| **EGCG** | Tocris | 4524 | 50 | DMSO |

### Proliferation assays

NSCs were plated 5000 cells per well in a 96 well plate and treated with pharmacological inhibitors or vehicle for 15 minutes prior to the addition of glutamate/aspartate or vehicle. Two days later, 20 µM 5-Ethynyl-2′-deoxyuridine (EdU; Click Chemistry Tools 1149) was added to the cell culture media for 30 minutes to label NSCs in S-phase. The cells were then fixed in 4% PFA for 20 minutes prior to undergoing a click reaction to detect EdU. Cells were permeabilized for 20 minutes in 0.5% Triton X-100 and incubated in the dark for 30 minutes with copper-dependent click reaction mixture (Click Chemistry Tools #1350) to label EdU directly with fluorophores (Azide-488 or –647). Cells were stored at 4 °C in the dark until imaging (see above). After imaging, a standard threshold was applied to the channel containing EdU staining for all images, and the analyze particles function was applied to count the cells with EdU signal above threshold. EdU counts are expressed as relative change to control within experiment to facilitate combining multiple experimental replicates. In generally, a minimum of 3 experimental replicates of each EdU proliferation assay was performed, with multiple well replicates per groups in each individual experiment.

### Glutamate clearance assay

Glutamate clearance experiments were performed on NSCs grown in 96 well plates to approximately 95% confluency. The growth media was removed and cells were rinsed twice with oxygenated artificial cerebrospinal fluid (aCSF) prepared according to the following recipe (in mM): 116 NaCl, 3.0 KCl, 1.25 NaH2PO4, 23 NaHCO3, 10 glucose, 2.0 MgSO4, and 2.0 CaCl_2_^53^. Cells were then pre-incubated for 15 minutes with aCSF containing EAAT inhibitors or vehicle before replacing the solution with 50 µL of aCSF containing 5 µM glutamate and EAAT inhibitors or vehicle. Immediately after (T0) and 5, 10, and 20 minutes later, samples (10 µL) were injected and glutamate quantified by uHPLC-ECD.

### uHPLC-ECD

Glutamate was quantified using uHPLC-ECD (ALEXYS Neurotransmitter Analyzer;Antec Scientific, USA). Samples were pre-column derivatized with o-phthaldialdehyde (OPA) and sulphite. Chromatographic separation was performed via an Acquity UPLC HSS T3 1.0 × 50 mm, 1.8 µm column (Waters).). The mobile phase A, for separation consisted of 50 mM phosphoric acid, 50 mM citric acid, 0.1 mM EDTA, and 1.5% v/v acetonitrile with a final pH of 3.1. Mobile Phase B, for post-separation flush, consisted of the same mobile phase A but with 50% v/v acetonitrile. The system was operated at a flow rate of 200 μl/min at a pressure of 380 bar. Electrochemical detection used a working potential of 850 mV (vs Ag/AgCl reference) and a range setting of 5 nA/V. Sample peaks were fitted to a glutamate standard curve.

### Plasmid design and lentiviral packaging

The CRISPRi lentivirus construct was modified from pLV hU6-sgRNA hUbC-dCas9-KRAB-T2a-GFP (Addgene #71237, a gift from Charles Gersbach^28^), which expresses all necessary CRISPRi machinery (both the dCas9-KRAB and sgRNA) from the same plasmid. The UbC promoter was replaced with an EF1α promoter to create pLV hU6-sgRNA EF1α-dCas9-KRAB-T2A-GFP. Additionally, two Esp3I recognition sites were placed after the U6 promoter for restriction cloning of sgRNA insert sequences. The insert sequences were synthesized as single strand oligonucleotides (Integrated DNA Technologies) in the form of 5’ GGACG(N)20 3’ and 5’ AAAC(N’)20C 3’ where (N)20 refers to the sequence of the sgRNA and (N’)20 is the reverse complement. These oligonucleotides were annealed together and ligated with Esp3I-digested pLV hU6-sgRNA EF1α-dCas9-KRAB-T2a-GFP to obtain the final constructs for lentivirus packaging. The sgRNA targeting the region -50 to +300 bp relative to the transcriptional start sequence of Slc1a3 were designed using the CRISPRi function of the online Broad Institute GPP portal^54,55^. The top ranked Slc1a3-targeting sgRNA returned by the GPP tool (5’CGGAGTAACAGCTTAGCGAG) and a non-targeting (NT, 5’GCGAGGTATTCGGCTCCGCG) sgRNA^56^ were used to create an EAAT1 KD plasmid and an NT control plasmid. These plasmids were packaged as VSV-G pseudotyped, second-generation lentivirus vectors (titers 1.51 × 10^9^ IFU/mL, EAAT1 KD virus; 1.53 × 10^9^ IFU/mL, NT control virus) by Vigene Biosciences, Inc. These plasmids will be made available via addgene upon publication.

### CRISPRi KD in vitro

Cultured adult DG NSCs were plated 5000 cells per well in 96 well plates and treated with EAAT1 KD or NT control virus (∼MOI 75 - 150). Cells were maintained in culture for 8 – 12 days after virus treatment and were passaged 1 – 2 times during that timeframe. Cells were treated with 100 µM glutamate or vehicle 2 days before a 30 minute pulse of 20 µM EdU and fixation. Immunofluorescence labeling, EdU detection, and imaging were performed as described above. ImageJ was used to manually draw ROIs around GFP+ and GFP-NSC cell bodies in the brightfield channel and measure the average EAAT1 immunofluorescence signal intensity within the ROI. The number of GFP+EdU+ cells was quantified by manually identifying all GFP+ cells and evaluating colocalization with EdU.

### Stereotaxic surgery

Mice were anesthetized by isoflurane inhalation (Akorn, 4% induction, 2% maintenance) in oxygen and mounted in the stereotaxic apparatus (Stoelting). The scalp was sterilized with alcohol (Fisher) and betadine (Fisher) before exposing the skull via a single scalpel blade incision. Bilateral bur holes were drilled into the skull, and Hamilton syringes were positioned at A/P -2.0 mm and M/L +2.0 mm before being slowly lowered to a depth of DV –1.9 mm (all coordinates relative to bregma). Mice were administered 0.9 μL EAAT1 KD virus in one hemisphere and 0.9 μL NT control virus in the contralateral hemisphere at a rate of 0.1 μL/min via an automated injector system (Stoelting). The brain hemisphere receiving EAAT1 KD virus was counterbalanced within each timepoint cohort and across sexes. Mice were administered carprofen prior to surgery and daily for 3 days post surgery as well as buprenorphine immediately post surgery. 36 mice total received lentiviral infusion. Cohort 1: 16 (8 male, 8 female) were given 3 injections of EdU (150 mg/kg, i.p.) 4 hours apart 1 week after lentiviral infusion then perfused. GFP expression, indicating proper infusion placement, was confirmed by a blinded observer in 5 NT and 10 EAAT1 KD DGs. Cohort 2: 10 (5 male and 5 female) received a single daily injection of BrdU (150 mg/kg, i.p.; Sigma B5002) on days 6 – 8 after infusion and 3 injections of EdU on day 21 after lentivirus injection. GFP expression was confirmed by a blinded observer in 8 NT and 8 EAAT1 KD DGs. Cohort 3: 10 mice (5 male and 5 female) received a single daily injection of BrdU (150 mg/kg, i.p.; Sigma B5002) 1 month after infusion, 1 mo before 3 EdU injections and perfusion. GFP expression was confirmed by a blinded observer in 8 NT and 6 EAAT1 KD DGs. All cohorts were perfused two hours after the final EdU injection.

### Experimental treatment and RNA extraction for RNAseq

NSCs of passage 4 were plated 75,000 cells per well in a 12 well plate and treated with 1 µM TFB-TBOA or vehicle for 15 minutes prior to the addition of 100 µM glutamate or vehicle (n = 3 wells per treatment). After 48 hours, cells were harvested with Accutase (StemCell technologies) and a subset (approximately 75,000 cells) was pelleted prior to RNA extraction using the NucleoSpin RNA Plus XS kit (Takara) following manufacturer protocol. Assessment of RNA quality with Qubit RNA HS assay kit (Invitrogen) revealed RIN values of at least 8 for all samples.

### RNAseq library generation, sequencing, alignment, and transcript quantification

Library generation was performed as previously described^27^ using the Clontech SMART-Seq HT (Takara) kit. Purified library products were then submitted to HiSeq 4000 paired-end sequencing (Illumina) with a depth of 15 – 20 million 2 × 150 bp clusters. AdapterRemovalv2.2.0 was used to trim individual FASTQ files for adapter sequences and filter for a minimum quality score of Q20. HISAT2 v2.0.6 was used to exclude reads aligning to a composite reference of rRNA, mtDNA, and PhiX bacteriophage sequences obtained from the NCBI RefSeq. Primary alignment was performed to the mouse reference genome GRCm38p4 using HISAT2. Gene expression values were quantified for all genes described by the GENCODE GeneTransfer Format (GTF) release M14 (mouse) using the featureCounts tool of the Subread package v1.5.1 58 in stranded mode. Raw and processed counts will be made available via GEO upon publication.

### Differentially expressed gene (DEG) analysis

DEG analysis was performed using pair-wise comparisons between the following treatment groups: vehicle/vehicle vs glutamate/vehicle, vehicle/vehicle vs vehicle/TFB-TBOA, glutamate/vehicle vs glutamate/TFB-TBOA, and vehicle/TFB-TBOA vs glutamate/TFB-TBOA. For all comparisons, the DESeq R package^57^ was used to compute log_2_fold changes in gene expression between treatment groups, and genes with a Benjamini-Hochberg adjusted p-value < 0.05 were considered as DEGs.

### Gene ontology enrichment (GO) analysis and clustering

GO analysis of DEGs was performed using the ClueGo plugin v.2.5.7^58^ for Cytoscape v3.9.0. Upregulated and downregulated DEGs were analyzed separately. DEGs were compared to the GO term annotations within the reference ontology GO_BiologicalProcess-EBI-UniProt-GOA-ACAP-ARAP_08.05.2020_00h00, and a two-sided hypergeometric test was performed with a Bonferroni adjusted p value <0.05 considered as significant enrichment. Similar GO terms were clustered using iterative kappa score grouping.

### Western Blot

NSCs were plated 200,000 cells per well in 6 well plates and treated with 1 µM TFB-TBOA or vehicle for 15 minutes prior to the addition of glutamate or vehicle. Two days later, cells were detached by scraping in ice-cold PBS and pelleted by centrifugation at 800 x g for 2 minutes. Pellets were resuspended, vortexed vigorously, and sonicated in ice-cold lysis buffer consisting of RIPA buffer (Fisher P189900), 1% Halt protease and phosphatase inhibitor (Fisher P178440), and 1% EDTA. Protein lysate concentrations were quantified using the Pierce Rapid Gold BCA protein assay kit (Fisher A53225).

Samples were prepared for SDS-PAGE by diluting 10 µg of protein lysate in Laemmli buffer (Biorad 1610747) with 5% β-mercaptoethanol. Samples were heated at 95°C for 10 min and loaded into a 4-15% gradient Tris-glycine gel (Biorad 4561084) immersed in Tris-glycine-SDS running buffer (Biorad 1610772). Samples were run for 1 hour at 120 V and then transferred overnight at 4°C with 30 V to a nitrocellulose membrane (Fisher NC9680617) in Tris-glycine-20% methanol transfer buffer. The membrane was rinsed with TBS, incubated for 1 hour in 5% BSA/TBS/0.1% Tween20 (blocking buffer), and incubated overnight at 4°C in mouse anti-mTOR (Cell Signaling 4517) and rabbit anti-phospho(p)-mTOR(Ser2448) (Cell Signaling 2971) both diluted 1:1000 in blocking buffer. Membranes were then rinsed 3x in TBS and incubated for 1 hour in anti-mouse IR680 (VWR 102971-020) and anti-rabbit IR800 (VWR 103011-494) both diluted 1:5000 in blocking buffer. Membranes were then rinsed 3x in TBS and imaged in the 680 and 800 channels using an Odyssey CLx (LI-COR).

Immunoreactive bands for mTOR and p-mTOR were analyzed using ImageStudio (LI-COR). A hand-drawn box of consistent size was traced over each band, and signal intensity was measured with background subtraction using the median of the region immediately surrounding each band. The ratio of p-mTOR to total mTOR signal was calculated for each band.

### Statistics

Statistical analysis was performed using GraphPad Prism version 9 or higher, except for RNAseq data, which is described above. Generally, if experiments used a 2×2 factor design, 2-way ANOVA was used with error-corrected posthoc tests. If experiments used a 3×2 factor design, 3-way ANOVA was used with error-corrected posthoc tests. If experiments only had 2 groups, non-parametric comparisons were generally used (Mann Whitney). Details of statistical tests are in Table S4.

## Acknowledgements

This research was supported by R21 NS123797 from NINDS/NIH to EDK.

Research reported in this publication was supported by The Ohio State University Comprehensive Cancer Center and the National Institutes of Health under grant number P30 CA016058. We thank the Genomics Shared Resource at The Ohio State University Comprehensive Cancer Center, Columbus, OH for RNAseq services. We also thank the Cancer Program IT resource for sponsoring site licenses for GraphPad Prism and Biorender.

## Author contributions

Conceptualization, JDR and EDK; Methodology, JDR and EDK; Investigation, JDR, IRB, JEC, EA, DE, AW, and VV; Writing – Original Draft, JDR and EDK; Writing – Reviewing and Editing, JDR and EDK; Supervision, JPB and EDK; Funding Acquisition, EDK.

## Supplementary Figures

**Fig S1: Supplement to Fig 1.**
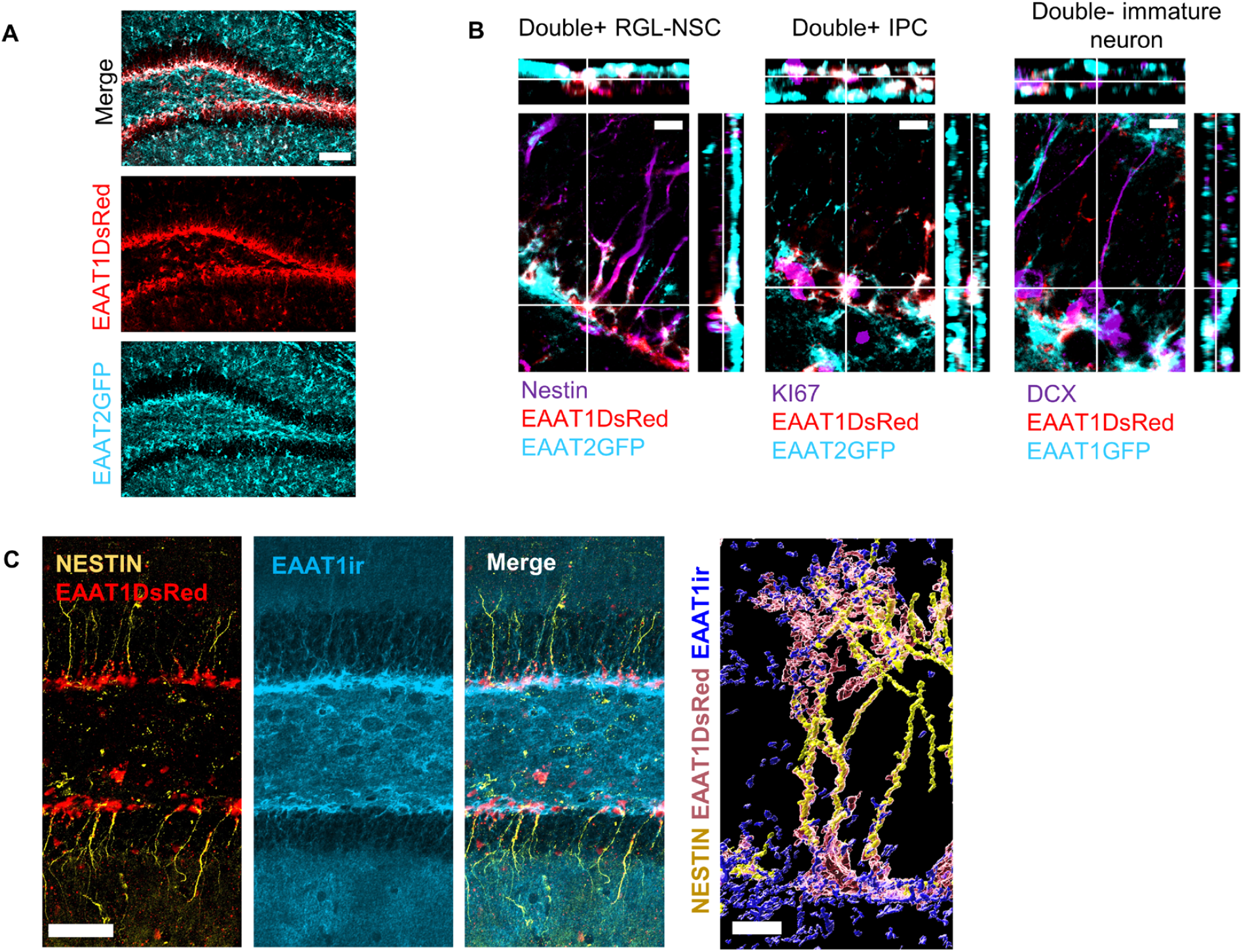
A) EAAT1DsRed+EAAT2GFP+ fluorescence in the adult DG. Scale = 100 µm. B) Orthogonal projections of example: EAAT1DsRed+EAAT2GFP+ (double+) Nestin+ radial glia like (RGL) NSC, EAAT1DsRed+EAAT2GFP+ (double+) Ki67+ IPC, EAAT1 DsRed-EAAT2GFP- (double-) DCX+ immature neuron. Scale = 10 µm. C) Nestin immunolabeling in DG of EAAT1DsRed+ co-localized with EAAT1 immunoreactivity (ir). Left is merged z-stack. Right is 3D reconstruction showing an example Nestin+EAAT1DsRed+ radial glia like NSC with EAAT1ir in the cell body and apical terminals.

**Fig S2: Supplement to Fig 2.**
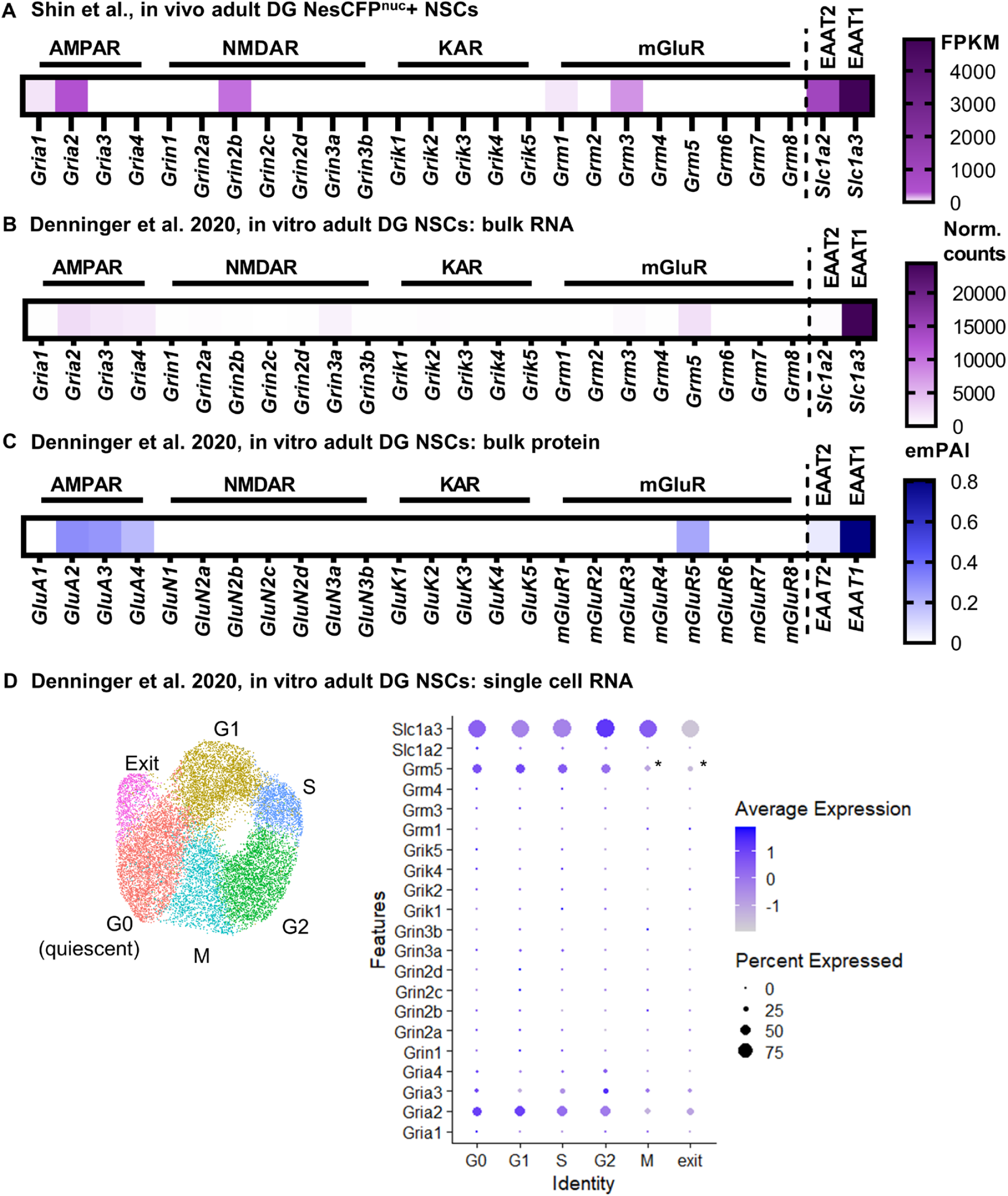
A) FPKM of glutamate receptor and EAAT1/2 genes from Shin et al., 2015. B) Transcript length-normalized RNAseq counts of glutamate receptor genes and EAAT1/2 from cultured adult DG NSCs in Denninger et al., 2020. C) emPAI length-normalized semi-quantitiative spectral counts for glutamate receptor and transporter proteins from cultured adult DG NSCs in Denninger et al., 2020. D) Single cell RNAseq of cultured DG NSCs from Denninger et al., 2020 reflects a mixture of active and quiescent NSCs plus a cycle-exiting population, as shown in UMAP. Dot plot shows average normalized RNA count per cell (color scale) and percent of cells expressing (dot size) glutamate transporter and receptor genes. * = differentially expressed gene for cluster in original analysis.

**Fig S3: Supplement to Fig 3.**
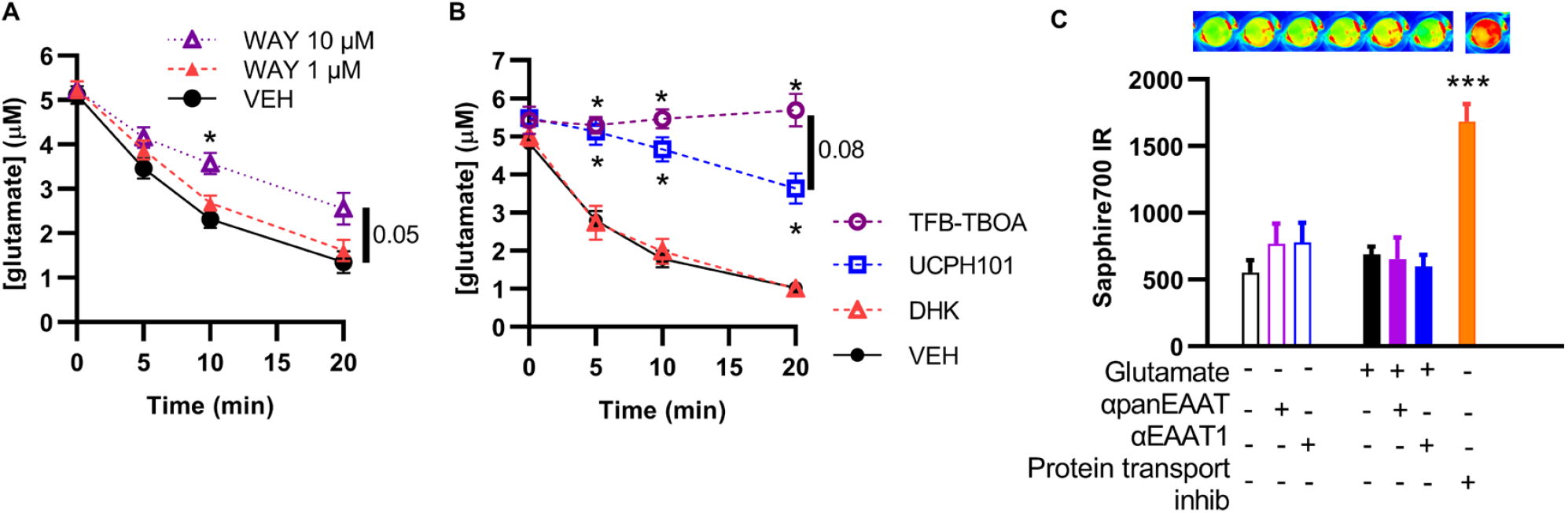
A) Glutamate in NSC conditioned media as measured by uHPLC after a 5 µM pulse after pre-treatment with an EAAT2-selective (1 µM) and an EAAT1 and2 selective (10 µM) dose of WAY213613. *p<0.05 vs 1 µM and vs 10 µM. Mean ± SEM of N = 6 independent experiments. B) Glutamate in NSC conditioned media as measured by uHPLC after a 5 µM pulse after pre-treatment with a panEAAT inhibitor (TFB-TBOA), an EAAT1 inhibitor (UCPH101), or and EAAT2 inhibitor (DHK). *p<0.05 Tukey’s comparisons within timepoint. Mean ± SEM of N = 3 independent experiments. C) Sapphire IR signal in cultured NSCs after treatment with glutamate +/− panEAAT inhibitor (TFB-TBOA) or EAAT1 inhibitor (UCPH101). Protein transport inhibitor at toxic dose is positive control for presence of apoptosis in Sapphire IR700 assay. ***p<0.001 Tukey’s multiple comparisons. Mean ± SEM of N = 3-6 replicates/experiment, 2 independent experiments. Top shows representative image of Sapphire IR signal in treatment groups.

**Fig S4: Supplement to Figure 4.**
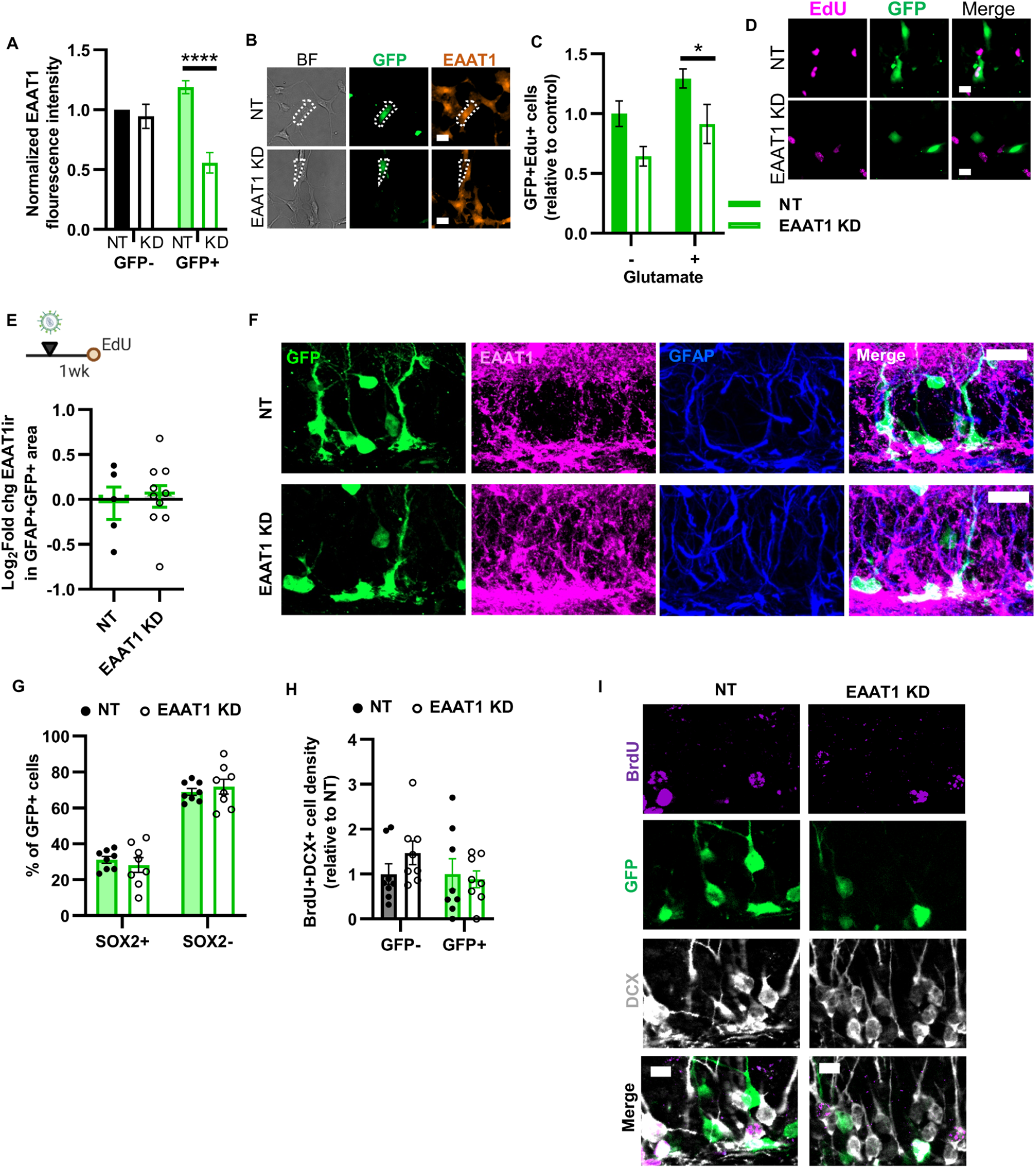
A) EAAT1 immunoreactive intensity in cultured NSCs infected with NT and EAAT1 KD CRISPRi lentiviral vectors. ****p<0.0001 Sidak’s multiple comparisons within GFP type. Mean ± SEM of N = 4 independent infections. B) Representative images of EAAT1 labeling coupled with GFP and brightfield to visualize all NSCs. Scale = 10 µm. C) Edll+GFP+ cell counts in cultured NSCs infected with NT and EAAT1 KD CRISPRi lentiviral vectors. Mean ± SEM of N = 3 replicates/infection, 3 independent infections. D) Representative images of EdU labeling coupled with GFP in NSCs infected with NT and EAAT1 KD CRISPRi lentiviral vectors. Scale = 10 µm. E) Timeline and EAAT1 immunolabeling intensity within GFP+GFAP+ area of mice infused with NT or EAAT1 KD CRISPRi lentiviral vectors in the DG then perfused 1 week later. Mean ± SEM of N = 5-8 individual mice (points shown). F) Representative image of EAAT1 labeling coupled with GFP and GFAP at 1 week after CRISPRi infusion. Co-labeling becomes white in merged image. Scale = 20 µm. G) Percent of GFP+ cells co-expressing SOX2 3 weeks after CRISPRi infusion. H) Density of GFP+ and GFP-BrdU+DCX+ immature neurons/neuroblasts 3 weeks after CRISPRi infusion. I) Representative image of BrdU, GFP, DCX co-labeling 3 weeks after CRISPRi vector infusion. Scale = 10 µm.

